# Anneal-MaxCal: kinetic ensembles from a Markov model learned along a solute-tempering ladder

**DOI:** 10.64898/2026.10.05.756723

**Authors:** Z. Faidon Brotzakis, Nikolaos Rekatsinas

## Abstract

A full characterisation of a molecular system requires its metastable states, their populations and their interconversion rates. Molecular Dynamics can in principle provide this characterisation but is limited by sampling and by force-field inaccuracies. Enhanced sampling addresses the first problem, but typically requires prior knowledge of reaction coordinates or metastable states and often sacrifices the kinetics, while maximum-caliber reweighting by Brotzakis, Vendruscolo and Bolhuis [Proc. Natl. Acad. Sci. U.S.A., 118(2):e2012423118, 2021] has been implemented to that matter on reweighting continuous pathways by imposing rate constants as kinetic constraints in two-state protein folding. Here, we introduce Anneal-MaxCal, which tackles both problems by imposing rate constants or time-resolved data on a Markov model learned along an annealed solute-tempering ladder spanning softer to physical interactions. MBAR assigns physical free energies to all 160 cells of a slow TICA subspace, against 79 visited at the physical coupling. The square-root approximation (SqRA), a discretised diffusion on this free-energy landscape, converts these populations into a detailed-balance rate matrix. Its prefactors, which act as a position-dependent diffusivity across each cell face, are learned from transitions at the softer couplings and extrapolated log-linearly to the physical coupling. This yields rates and a committor across the TICA space, although the physical rung contains a single barrier crossing. For alanine dipeptide, 665 ns of simulation give a relaxation time of 5430 *±* 996 ps against 4413 *±* 446 ps from 15.4 *µ*s of unbiased dynamics, whereas a constant prefactor is 2.5 times too slow. Maximum caliber then minimally perturbs this prior, with the posterior populations following from a generalised Doob transform. A synthetic time-resolved density with a tenfold slower back-transition lowers Δ*F* from 5.191 to 2.885 *k*_B_*T*, raises the product-basin population from 0.0055 to 0.0529 and slows relaxation 9.47-fold, leaving the reactant basin unchanged. Forward and backward rates accelerated 3- and 20-fold raise Δ*F* by +1.892 *k*_B_*T* and shorten relaxation from 4818 to 289 ps.

## 1 Introduction

From the point of view of statistical mechanics, a full characterisation of a molecular system requires its metastable states, their equilibrium populations and the rates of interconversion between them. Molecular dynamics (MD) simulations can in principle provide all three, but in practice face two limitations. The first is the rare-event problem: when free-energy barriers are large compared with *k*_B_*T*, transitions between states are too infrequent to be sampled directly. The second is the accuracy of the force field, whose approximations propagate into both the thermodynamics and the kinetics, and which affect rates particularly strongly, since these depend exponentially on barrier heights.

Enhanced-sampling methods address the first limitation in several ways. Tempering methods raise the temperature or scale the Hamiltonian [37, 26]. Potential-biasing methods add a bias along chosen collective variables [38, 39]. Path-based methods harvest reactive trajectories between predefined states [40, 41, 42]. Adaptive sampling methods launch short unbiased trajectories guided by a Markov state model (MSM) [43, 44]. These methods typically require prior knowledge of reaction coordinates, collective variables or metastable states. Tempering and potential-biasing methods, moreover, recover states and populations at the expense of the kinetics, which must then be reconstructed by dynamical reweighting [7, 8, 9]. MSMs assemble many short trajectories into long-time kinetics [1, 2], but their accuracy is limited by the equilibrium weight assigned to the trajectory starting points, which short simulations launched away from equilibrium bias [4]. Multi-ensemble Markov models and dynamic histogram analysis combine simulations from several thermodynamic states into a single kinetic model [5, 6].

The second limitation has been addressed by incorporating experimental data into simulations through the maximum entropy and maximum caliber principles [13, 14, 15]. Maximum caliber has been used to correct MSMs towards measured kinetics [18], and to impose rate constants as kinetic constraints on path ensembles from MD and transition path sampling, a method termed continuum path ensemble maximum caliber (CoPE-MaxCal) [19, 31], which has been extended to optimise force-field parameters against kinetic data [20]. Time-resolved data have been imposed through replica-averaged restrained simulations [21] and, more recently, through time-resolved Bayesian/maximum entropy (trBME) refinement [45], which combines a Fokker–Planck model of the dynamics along a chosen coordinate with structural ensembles from simulation. Related approaches reconcile MSMs with equilibrium [22] and dynamic [23] experimental observables. Most of these methods, however, require a reaction coordinate or predefined states: CoPE-MaxCal reweights a prior path ensemble between two predefined states along a chosen collective variable, and trBME propagates the dynamics along a coordinate that must be specified in advance.

Here, we introduce Anneal-MaxCal, which addresses both limitations within a single framework. Walkers are annealed along a solute-tempering ladder [26, 10, 11], from a softened Hamiltonian at which barrier crossings are frequent to the physical one. MBAR [12] provides the equilibrium populations at every coupling, including in regions that the physical coupling never visits. From these populations, the square-root approximation (SqRA) [3] builds a rate matrix that obeys detailed balance. Its kinetic prefactors, which act as a position-dependent diffusivity, are learned from transitions at the softer couplings and transferred to the physical coupling. Transition counts at the physical coupling are used only for validation. No predefined reaction coordinate is required: the dynamics is propagated by the full Markov model over a generic slow subspace, and the committor that indexes the subsequent reweighting is computed on the learned model. This prior model is then minimally perturbed by maximum caliber to reproduce kinetic data, either committed rates or time-resolved densities. The posterior is obtained by a generalised Doob transform [16, 17], so that the posterior stationary distribution is derived rather than fitted, and purely kinetic constraints yield thermodynamic predictions.

We apply Anneal-MaxCal to alanine dipeptide in vacuum, for which 15.4 *µ*s of unbiased MD provide an independent reference for both populations and rates. We then impose two types of synthetic kinetic data, for which the true correction is known: a time-resolved density and committed forward and backward rates. The method is assembled and validated one stage at a time as ladder, slow subspace, discretisation, MBAR, generator and maximum caliber — with an independent check at each stage. This identifies both the conditions under which the approach holds and its remaining limitations.

## 2 Theory

### 2.1 System and conditions

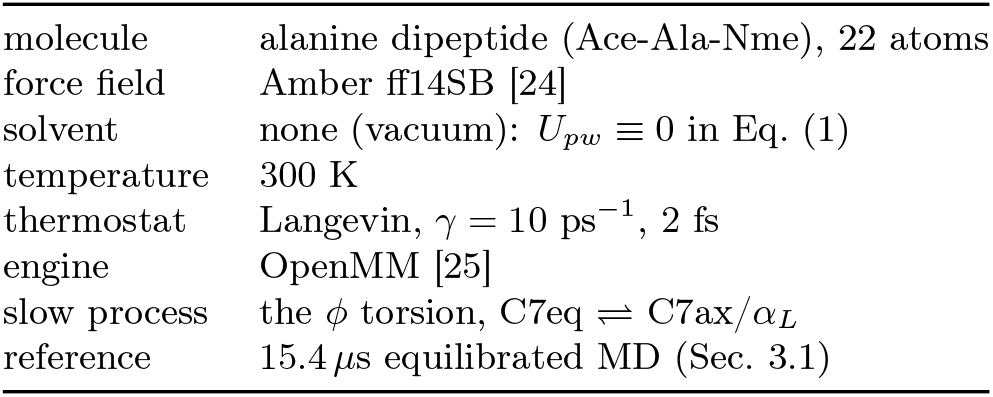

|  |  |
| --- | --- |
| molecule | alanine dipeptide (Ace-Ala-Nme), 22 atoms |
| force field | Amber ff14SB [24] |
| solvent | none (vacuum): $U_{pw} \equiv 0$ in Eq. (1) |
| temperature | 300 K |
| thermostat | Langevin, $\gamma = 10$ ps$^{-1}$, 2 fs |
| engine | OpenMM [25] |
| slow process | the $\phi$ torsion, $C7eq \rightleftharpoons C7ax/\alpha_L$ |
| reference | 15.4 μs equilibrated MD (Sec. 3.1) |

### 2.2 Notation

Throughout, *x* is a configuration of the 22 atoms, *β* = 1*/k*_B_*T*, and *U*_*λ*_ is the potential at coupling *λ* (Sec. 2.3). Continuum densities over configuration space are written *P*, with a subscript naming the ensemble: 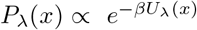 is the Boltzmann density at coupling *λ*, and *P*_1_ the physical one. Transition densities are written *ρ* with a bar, *ρ*_*λ*_(*x*’|*x*). Discrete objects on the *n* cells carry Latin indices: *T*_*ij*_ is a transition matrix, *π*_*i*_ its stationary vector, *C*_*ij*_ transition counts, *Q*_*ij*_ a rate matrix (generator). The symbol *π* is reserved for the discrete stationary vector and is related to the continuum only through Eq. (10). Time-resolved observables are written *ρ*^*\**^(*ϕ, t*) for data and *ρ*_*µ*_(*ϕ, t*) for the model. Tildes denote posterior objects. Free energies are in units of *k*_B_*T*.

### 2.3 The annealed ladder

#### Targets

Direct sampling of *P*_1_ is barrier-limited, so we interpolate towards it from a softened ensemble. Under the solute-scaling (SST2/REST2) partition [26] the potential splits into a part that is never scaled, *U*_hard_ (bonded terms, and solvent–solvent interactions when the solvent is explicit), the solute–solute part *U*_*pp*_ (torsions and intramolecular nonbonded terms) scaled by *λ*, and the solute–solvent part *U*_*pw*_ scaled by 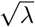,

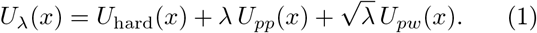

Everything below is written for this general form. Its only requirement is that every stored configuration carries the two scalars *U*_*pp*_(*x*) and *U*_*pw*_(*x*), because the potential at *any* coupling is then known for that configuration. *U*_hard_ is common to all couplings and cancels. In implicit solvent *U*_*pw*_ is the generalised-Born solvation energy, while with explicit solvent it is the solute–solvent interaction. **Alanine dipeptide in vacuum, the system studied here, is the special case** *U*_*pw*_ *≡* 0, for which *U*_*λ*_ is linear in *λ*. The ladder is an increasing sequence *λ*_0_ *< λ*_1_ *< · · · < λ*_*K*_ = 1.

#### The softest rung is one continuous trajectory

The information a ladder carries about the barrier is set by how often the system crosses it, and crossings per unit time are fixed by the coupling, not by how the time is spent. At the softest rung that time is therefore spent on a single continuous equilibrium trajectory rather than on short walkers, so that every crossing is a genuine equilibrium crossing and lags up to nanoseconds are available there. The coupling itself is chosen by measurement (Sec. 3.2): softening raises the crossing rate roughly exponentially while degrading the overlap with the next rung, and *λ*_0_ = 0.20 is the softest value from which a single free-energy-perturbation step still reaches the rest of the ladder. It is run for 10 ns, starting from the extended alanine-dipeptide structure of the openmmtools test system (*ϕ* = *ψ* = 180°, on the reactant side of the barrier) after 200 ps of discarded equilibration at *λ*_0_. At this coupling the run makes 72 committed crossings in 10 ns, so the starting basin is forgotten within the first nanosecond.

#### Mutation and selection

Each subsequent rung applies two operations to an ensemble of *M* = 2048 walkers, named in the standard sequential Monte Carlo vocabulary [10, 11]. *Mutation* is molecular dynamics: the one-step transition density *ρ*_*λ*_(*x*’ | *x*) of the Langevin integrator leaves its Boltzmann density invariant,

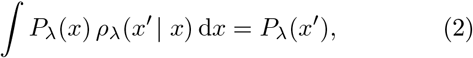

exactly for the continuous dynamics and to *O*(Δ*t*^*p*^) for a finite step, and so does the *τ*_r_ = 20 ps segment that is its convolution. *Selection* is by the free-energy-perturbation (FEP) factor. Moving the target from *λ*_*k*_ to *λ*_*k*+1_ changes the potential of a fixed configuration by

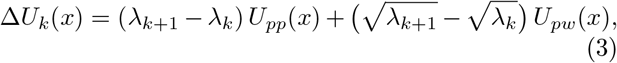

and the corresponding weight is the Zwanzig exponential,

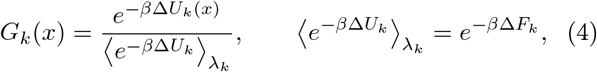

so the step that transports the ensemble and the step that estimates the free energy are the same object. Because Δ*U*_*k*_ depends on *x* alone and the dynamics at rung *k* is run at its own target, the path density cancels from every importance ratio: no Girsanov factor, no Jarzynski work [27] and no accumulated action appears [7, 8].

#### Protocol

The ladder is *threaded* from the continuous run. The candidates for the first walker rung are *all* stored frames of the *λ*_0_ trajectory, each an equilibrium sample of 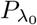. They are weighted by Eq. (4) and *M* initial conditions are drawn by systematic resampling [28]. Every later rung then (i) chooses *λ*_*k*+1_ by the step policy of Sec. 2.4 from the walkers’ endpoint energies, at no simulation cost, (ii) reweights the endpoints, ln 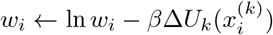, (iii) resamples only if the accumulated effective sample size has fallen below 0.5, and (iv) propagates *τ*_r_ = 20 ps at *λ*_*k*+1_, storing a frame every 0.2 ps. Only positions are inherited: velocities are redrawn from the Maxwell– Boltzmann distribution at every rung, which samples the exact conditional because momenta are independent of configuration in the Boltzmann measure. Every stored frame, not only the endpoints, enters the reweighting of Sec. 2.5.

### 2.4 Adaptive coupling selection and walker weights

The weight *w*_*i*_ belongs to walker *i* and is a configuration-space object, the density ratio 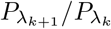 at its endpoint, so that ∑_*i*_ *w*_*i*_*f* (*x*_*i*_)*/ ∑*_*i*_ *w*_*i*_ estimates 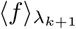. It is reset at every resampling and is an internal device of the sampler: the weights used for every reported estimate are the MBAR weights of Sec. 2.5, recomputed from scratch over all stored frames.

The step is not fixed in advance. With the normalised effective sample size

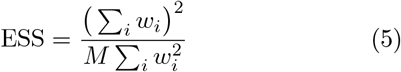

evaluated on the single-step weights exp[*− β*Δ*U* (*x*; *λ*_*k*_ *→ λ*’)] of Eq. (3), *λ*_*k*+1_ is located by 40 bisections so that this *incremental* ESS equals 0.8 [29]. For the first step the same rule is applied over all frames of the continuous run. Evaluated on the weights accumulated since the last selection, Eq. (5) is the *accumulated* ESS, which decays geometrically and alone triggers resampling, at 0.5.

#### What resampling does to a rare basin

Resampling is unbiased in expectation, but not in every realisation, and the distinction matters for a basin that holds a few walkers. The product basin holds ~ 1% of *P*_*λ*_, about 20 of 2048 walkers, and a 20 ps segment at stiff coupling never crosses back, so a single resampling step fixes the product-basin population of every later rung at whatever that draw produced. MBAR treats those rungs as equilibrium samples of their couplings. We therefore thread the ladder 2 times from the same continuous run, with independent resampling seeds, and pool the threadings in one MBAR problem. The effect of one, two and four threadings on accuracy and cost is measured in Secs. 3.3 and 3.6.

### 2.5 MBAR across the rungs

Each stored configuration *n* carries *U*_*pp*_(*n*) and *U*_*pw*_(*n*), so its reduced potential is available at *every* coupling,

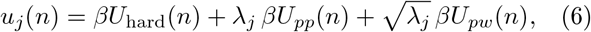

the configuration-dependent but coupling-independent *βU*_hard_ cancelling in every expression below. With *N*_*j*_ frames collected at *λ*_*j*_, the multistate Bennett acceptance-ratio (MBAR) equations [12, 30] determine the dimensionless free energies *f*_*j*_ self-consistently,

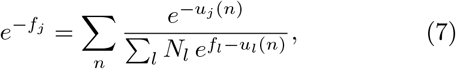

and the weight of every configuration at any coupling follows in one pass. At the physical Hamiltonian it is

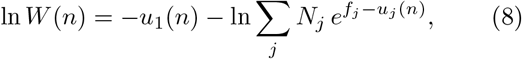

and at any other coupling by the exact re-tilt

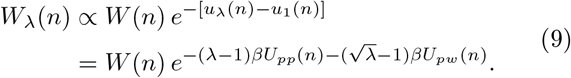

No binning in any coordinate is involved. The stationary vector of a discretisation into cells *S*_1_, …, *S*_*n*_ is obtained by summing within each cell,

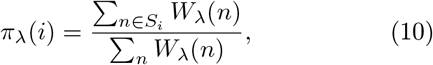

the sole bridge between the continuum and the discrete chain. This is the factor that carries the barrier: it is built from the *energies* of frames, including frames that never cross it.

### 2.6 Markov state models

#### Standard construction

For a partition into cells the transition matrix at lag *τ* is *T*_*ij*_(*τ*) = ℙ (*x*_*t*+*τ*_ *∈ S*_*j*_ | *x*_*t*_ *∈ S*_*i*_) [1, 2]. Distributions propagate by matrix powers, *π*^*T*^*T* = *π*^*T*^, and the relaxation timescales follow from the eigenvalues 1 *> µ*_2_ *≥ µ*_3_ *≥* … as

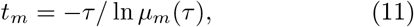

independent of *τ* exactly when the chain is Markovian at that lag. *t*_2_ is the barrier-crossing timescale used throughout. Microscopic reversibility imposes detailed balance, *π*_*i*_*T*_*ij*_ = *π*_*j*_*T*_*ji*_. When a chain is instead written as a rate matrix *Q* with *T* (*τ*) = *e*^*Qτ*^, its timescales are *t*_*m*_ = *−* 1*/v*_*m*_ from the eigenvalues *v*_*m*_ of *Q*, lag-independent by construction.

#### Committor

With *A* and *B* disjoint sets of cells for reactant and product and ℐ the intermediate cells, the forward committor *q*_*i*_ is the probability of reaching *B* before *A* from cell *i* [32, 33, *34]*,

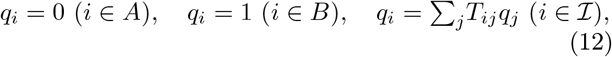

solved as (⊮ *− T*_*ℐℐ*_) *q*_*ℐ*_ = *T*_*ℐB*_**1**. *A* and *B* are the cells whose mean *ϕ* lies in the C7eq (*ϕ < −* 1) and C7ax/*α*_*L*_ (0.4 *< ϕ <* 2.2) cores.

### 2.7 Slow subspace and cells

#### Features and subspace

Frames are featurised generically, as sin and cos of the backbone torsions, and the slow subspace is found by time-lagged independent component analysis (TICA) [35, 36] at a lag of 2 ps. The basis is *linear* on purpose. A nonlinear slow coordinate is a better slow mode but saturates inside each basin, and a uniform tiling of a saturating coordinate collapses almost all density into a few cells with unbounded free-energy steps between the rest.

The basis is estimated at *λ* = 0.41, not at the softest rung. At *λ*_0_ the product basin is shallow enough that the slowest linear mode is dominated by fast escapes from it and by wide soft excursions, not by the barrier geometry that matters at *λ* = 1. How many cells then land on the barrier is measured in Sec. 3.2. The continuous *λ*_0_ run keeps its other role: a direct Markov model on it, Eq. (11), supplies the internal reference timescale against which the fit is checked, and no unbiased trajectory at *λ* = 1 enters the coefficient fit.

#### Cells

The leading two components are tiled by a uniform 16 *×* 16 lattice. Cells are instantiated where any rung has a frame, restricted to the largest connected component, with nearest-neighbour lattice adjacency. The tiling is *uniform*, not clustered: clustering allocates cells by density, and a barrier is where the density is not.

### 2.8 The *λ*-conditioned generator

The prior chain is a rate matrix on those cells, conditioned on the coupling,

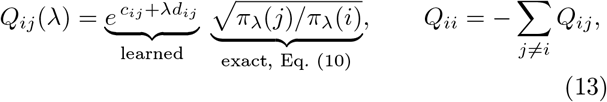

The second factor is the square-root approximation (SqRA) of a diffusive generator [3]. The use of a symmetric prefactor preserves the directional rate ratio imposed by the square-root factor, *Q*_*ij*_*/Q*_*ji*_ = *π*_*λ*_(*j*)*/π*_*λ*_(*i*), while allowing the kinetic scale of each neighbouring pair to be learned as a function of the coupling. The first factor is what the counts must supply, and *c*_*ij*_ and *d*_*ij*_ are its only parameters. For each pair of cells *i, j* that share a face of the lattice, 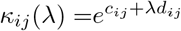 is the *kinetic prefactor* of the jump, in ps^*−*1^, which in the SqRA is proportional to the local diffusivity across the face. The corresponding reversible edge *conductance* is 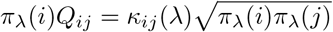. Thus,

- *c*_*ij*_ is its logarithm at *λ* = 0, the intercept.
- *d*_*ij*_ is the rate at which that logarithm changes with the coupling, the slope, so that ln *κ*_*ij*_ is linear in *λ*.

Both are symmetric, *c*_*ij*_ = *c*_*ji*_ and *d*_*ij*_ = *d*_*ji*_, which is what makes *π*_*λ*_(*i*)*Q*_*ij*_ = *π*_*λ*_(*j*)*Q*_*ji*_ hold, and both are defined only on the lattice: *Q*_*ij*_ = 0 unless *i* and *j* are neighbours. On the 16 *×* 16 lattice this is 288 faces, so 576 numbers in total, shared by all couplings, with no per-coupling or per-cell parameter. *c* and *d* do not change the imposed stationary populations, so what transfers from the soft couplings to *λ* = 1 is how fast the system moves between neighbouring cells for the supplied stationary populations, with unresolved barrier effects included in the kinetic prefactor.

The exponential ensures positivity, while the affine logarithm is a parsimonious first-order model of the coupling dependence. Log-linearity is a modelling assumption, rather than a consequence of detailed balance or of the affine vacuum potential. If an Arrhenius-type representation is introduced, 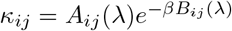, the assumption concerns the combined quantity ln *A*_*ij*_(*λ*) *− βB*_*ij*_(*λ*), with rates expressed numerically in ps^*−*1^. It does not require *A*_*ij*_ to be independent of *λ*, and *d*_*ij*_ is not generally a barrier derivative alone. Here *B*_*ij*_ is an effective barrier of the symmetric prefactor, rather than necessarily the directional activation barrier. Neither *A*_*ij*_ nor *B*_*ij*_ is separately fitted. Curvature in this combined dependence can limit transfer to couplings beyond those informing the fit (Sec. 4).

#### Imposed thermodynamics and learned kinetics

In the untruncated model, detailed balance with respect to the supplied *π*_*λ*_ holds at every coupling by construction, *π*_*λ*_(*i*)*Q*_*ij*_ = *π*_*λ*_(*j*)*Q*_*ji*_, so the equilibrium thermodynamics represented by these stationary populations is imposed, while the kinetic prefactor is learned. Transport to *λ* = 1 is therefore two different operations: the stationary populations are supplied through *π*_1_, while the kinetic prefactor is extrapolated log-linearly from the training couplings. On edges whose crossings are never observed the penalty on *d* pulls the extrapolation towards a *λ*-independent effective diffusivity.

#### Transition operator and physical time

Equation (13) is read directly: nothing is sampled, rolled out or decoded. Physical time is built in because the fit is to counts at known lags, and *T* and *π* are exactly what the maximum-caliber step consumes.

#### Training

Both *c* and *d* are fitted jointly by differentiation through the matrix exponential. Only *d* is explicitly penalised. One generator is fitted to four lags at once,

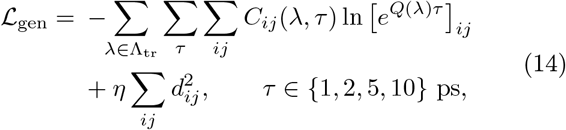

so that consistency under composition is structural rather than diagnostic. Counts are formed only from pairs *within* a trajectory: along the continuous run at *λ*_0_, and along each walker elsewhere. Training uses every coupling below 1, while **the physical rung’s transition counts are reserved for validation**, i.e. regulariser selection and short-time checks, and its configurations enter through *π*_1_. Symmetry is imposed at every step, 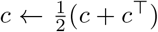 and likewise *d*. The half log-ratio of *π* is clamped to *±* 12 and the exponent *c* + *λd* to [*−* 15, 8], which prevents an unbounded rate across a large free-energy step. The matrix exponentials are formed in one batched scalin-and-squaring evaluation with the gradient through its adjoint. Optimisation is Adam at learning rate 0.05 for 3000 iterations from *c ~ −*3 + 0.1 *N* (0, 1), *d ~ N* 0.1 (0, 1), over four seeds. The regulariser *η* is selected by likelihood on the validation *λ* = 1 counts and never by reference to any timescale. The prior Markov chain is then 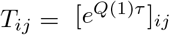 at *τ* = 20 ps, and the committor of Eq. (12) is computed on it.

### 2.9 Maximum-caliber reweighting

#### Where the committor enters

The committor *q*_*i*_ of Eq. (12) is computed once, on the prior chain, with *A* and *B* the torsional cores. It is the reaction coordinate of the reweighting, and it enters in three places. (i) It indexes the multiplier fields: the occupation field is expanded over (cell-mean *ϕ, q*) and the crossing field over *q* alone. (ii) It defines the crossing functional: a crossing is the passage of a committor level *q*_*k*_, so the crossing part of the tilt acts on transitions *along q* and on nothing orthogonal to it. (iii) It defines the rate observables, through the reactive flux and the committor-weighted populations of Eq. (20). There the committor of the *posterior* chain is used, recomputed at every evaluation, because a rate is a property of the chain being scored. The fields keep the prior’s *q*, so the parametrisation of the tilt does not move during the fit. Occupation alone cannot change a rate — a basin-to-basin transition completed within one lag never occupies a transition cell at a chain node — which is why the crossing field is needed.

#### The caliber tilt

Maximum caliber [13, 14, 15] makes the posterior path ensemble the prior times an exponential tilt in the constrained path functionals. On a Markov chain the two canonical functionals are the time spent in a cell and the unsigned number of crossings of a committor level — occupation and crossing activity [17]. With *ζ*_*i*_ the occupation multiplier of cell *i, α*_*k*_ the crossing multiplier of level *q*_*k*_ (14 levels) and 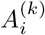 the indicator that cell *i* lies at or above committor level *q*_*k*_,

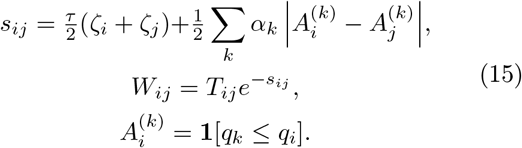

#### The multiplier basis

*ζ* is expanded in 10 Gaussian bumps over cell-mean *ϕ* and 9 over *q, α* in 9 bumps over *q*: 28 raw functions. These are not used as coordinates. The occupation bumps are centred, ∑_*i*_ *π*_*i*_*ζ*_*i*_ = 0, and orthonormalised under ⟨*f, g⟩* = ∑_*i*_ *π*_*i*_*f*_*i*_*g*_*i*_, while the crossing bumps are orthonormalised under the equilibrium flux across each level. Directions whose weighted norm falls below 10^*−*4^ of the largest are discarded, which leaves 6 occupation and 9 crossing directions. The coefficients of these 15 directions form *θ*. The reason is conditioning: a raw bump supported where *π* is 10^*−*7^ costs almost no caliber and runs away, and bumps over *ϕ* and over *q* acting on the same basin cells form a flat valley (Sec. 3.7).

#### Posterior construction via the generalised Doob transform

*W* is not stochastic, and row-normalising it would not give the maximum-caliber posterior. With (Λ, *ψ*) its leading eigenpair, *Wψ* = Λ*ψ*, the posterior chain [16, 17] is

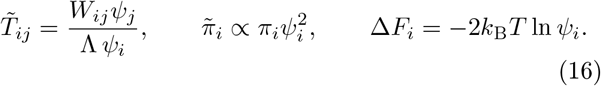

*s*_*ij*_ is symmetric, so *W* inherits the reversibility of *T* and (Λ, *ψ*) follows from a symmetric eigendecomposition. **The posterior stationary distribution is not fitted**: it falls out of *ψ*, which is what lets a purely kinetic constraint carry thermodynamic content.

#### How the time-resolved density is modelled

The model density is the posterior chain propagated from the prepared state *p*_0_, the Boltzmann-restricted product basin, and binned in *ϕ*:

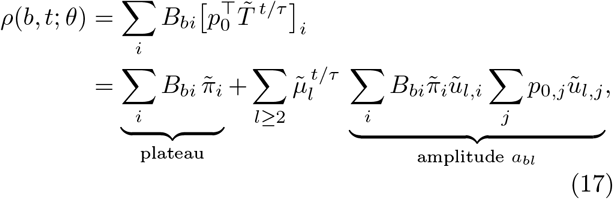

with *B*_*bi*_ = 1 when the cell-mean *ϕ* of cell *i* falls in bin *b*, and 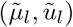 the eigenpairs of 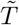, orthonormal under 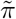. Every term is a function of *θ* only through 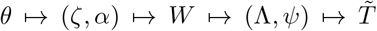. The equation separates what the data determine. The *long-horizon* density, 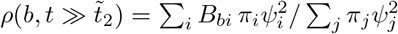 depends on *θ* only through *ψ*, that is only through Δ*F*_*i*_: the late checkpoints constrain the thermodynamics. The *approach* to it is set by the relaxation times 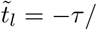 ln 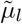 and by the amplitudes *a*_*bl*_: the early checkpoints constrain the kinetics. The 14 checkpoints are spaced geometrically from one lag to 8 times the target’s *t*_2_, so the plateau lies inside the fitted window. Nothing beyond it is fitted, and later times are read out by the same equation.

#### Objective

The multipliers minimise a misfit in the logarithm of the observables plus the caliber,

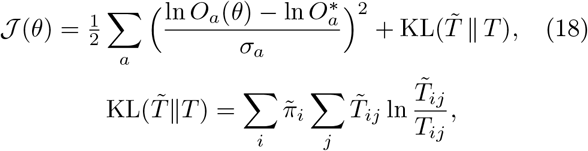

with the caliber the only regulariser [18, 19] and no Δ*F* penalty. *σ*_*a*_ is the standard deviation of the measurement error of observable *a* in ln *O*_*a*_, i.e. its relative uncertainty: *σ*_*a*_ = 0.15 means the observable is known to 15%, and it is what weighs each datum against the caliber. The data term is a Gaussian log-likelihood, so *J* is a negative log-posterior with the caliber as the prior.

#### Against a time-resolved density

The observables are the 12 *×* 14 entries of Eq. (17),

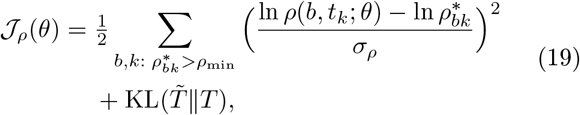

with *σ*_*ρ*_ = 0.15 and *ρ*_min_ = 10^*−*4^ the detection floor: populations no measurement could see do not enter.

#### Against rates

Rates replace the 168 transient observables by stationary ones. We use the committed rates of transition-path theory [33] of the posterior chain,

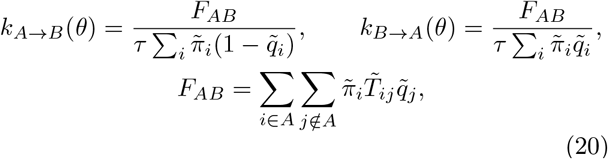

where 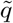 is the committor of 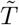 and *F*_*AB*_ the reactive flux. Their sum is the phenomenological relaxation rate, *k*_*A→B*_+ *k*_*B→A*_ *≈* 1*/t*_2_, and detailed bal ance ties their ratio to the equilibrium, 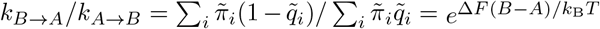. A *pair* of rates therefore fixes Δ*F*, while a single rate carries no information about it. The objectives are

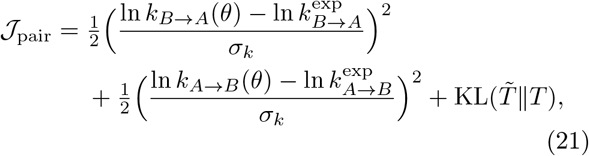

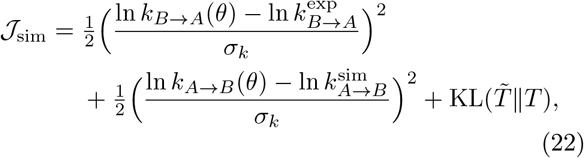

with *σ*_*k*_ = 0.05 and 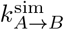 the prior chain’s own rate.

Compared with Eq. (19), three things change and nothing else does. The data are two stationary numbers rather than a transient, so no initial state, propagation or detection floor is involved. The committor, which in Eq. (19) only parametrises the tilt, now also defines the observable. And with 2 data for 15 multipliers the constraint is degenerate, so the caliber alone chooses among the chains that meet it. The tilt, basis, Doob map and optimiser are identical.

*J*_sim_ is *J*_pair_ with the unmeasured rate replaced by the simulation’s. This is an assumption, not data — that the simulation’s error lies in the measured direction — and it has a closed form, 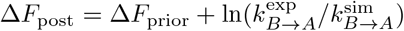. It is given the same width as the measured rate, so the fit meets it as it meets a measurement. The error it carries is therefore not a fitting weight but the error of the simulated rate, 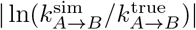, which passes one to one into Δ*F*

(Sec. 3.8). A rate is representable at lag *τ* only while *kτ «* 1, which caps speed-ups (though not slow-downs) and sets the lag.

#### Optimisation of the caliber multipliers

Every evaluation of *J* is a closed-form map

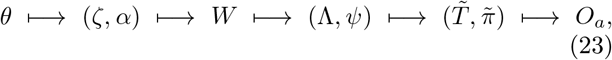

so no inner self-consistency loop is nested inside the fit. For the density it costs two symmetric eigendecompositions — one of *W*, one of 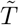, after which each checkpoint in Eq. (17) is a power of the stored spectrum. For rates, it costs one eigendecomposition and one linear solve for 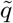. The optimisation is over the 15 coefficients and is *derivative-free*, because the map passes through an eigen-decomposition whose gradient is available but poorly conditioned near degeneracies. From each of two starting points — *θ* = 0, the prior, at which 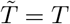 and *J* is the pure data misfit, and *θ* = 0.05 **1** — Nelder–Mead with adaptive simplex scaling is run to an absolute tolerance of 10^*−*10^ on *J*. Each result is polished by Powell’s direction-set method to a relative tolerance of 10^*−*12^, which removes the residual simplex-collapse error in the flat directions. The run with the lower *J* is kept. The search returns a large penalty when an eigendecomposition fails. With the orthonormal basis no bound on *θ* is active. The converged *θ*^*\**^ is not itself the deliverable and need not be unique. What is used is what Eq. (16) returns at 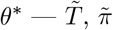 and Δ*F*_*i*_ — and convergence is judged on those.

#### Read-outs

The deliverables are 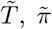 and Δ*F*_*i*_. Because the posterior is a reweighting of the ladder’s own frames,

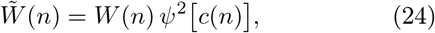

with *c*(*n*) the cell of frame *n*, free-energy surfaces follow per frame in any coordinates, and real configurations of the posterior are the stored frames themselves. Δ*F*_*i*_ is reported against cell-mean *ϕ* and against the committor *q*_*i*_, together with a kernel-smoothed representative 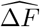 drawn only where cells exist within one bandwidth. Neither *F* nor Δ*F* enters any objective, so both are predictions.

## 3 Results

The construction was assembled one stage at a time, each checked against something independent of the next stage. Three stages failed on first construction. The failures and the changes they forced are reported where they occurred, because each locates a limit of the method.

### 3.1 Unbiased molecular-dynamics reference

The reference is unbiased molecular dynamics of the same system with the same integrator: 512 independent trajectories of 60 ns, launched with half the walkers in the C7eq basin (*ϕ ≈ −* 2.2, *ψ ≈* 2.1 rad) and half in C7ax/*α*_*L*_ (*ϕ ≈* 1.1, *ψ ≈ −*1.4 rad), so that *P* (*ϕ >* 0) = 0.5 at *t* = 0. That population relaxes to *≈* 0.01 within ~ 30 ns, and the first ~ 30 ns of every trajectory are discarded, leaving 512 *×* 30 ns = 15.4 *µ*s of equilibrated sampling, which gives *t*_2_ = 4413 *±* 446 ps and Δ*F* (*B − A*) = 4.747 *±* 0.205 *k*_B_*T* (*P* (*ϕ >* 0) = 0.0095). The reference enters no stage of the construction and is used only to judge it. How fast it converges is measured in Sec. 3.6.

### 3.2 Barrier sampling along the ladder

#### Choosing the softest coupling

A first continuous run at *λ* = 0.385, the softest rung of the walker ladder, crossed the barrier twice in 4.1 ns. Short scans at softer couplings gave 0.5, 1, 5 and 38 committed crossings per ns at *λ* = 0.385, 0.30, 0.20 and 0.10, while the FEP overlap with *λ* = 0.385 fell as 1, 0.79, 0.30 and 0.03. *λ*_0_ = 0.20 is the softest coupling from which one FEP step still reaches the ladder, and 10 ns there give 72 committed crossings. From it the ESS policy chose 8 further rungs, *λ* = 0.257, 0.327, 0.410, 0.509, 0.627, 0.767, 0.921, 1.000, with every incremental ESS at 0.8.

#### Barrier information

Pooled over the 2 threadings the ladder contains 449 committed crossings (Fig. 1a), essentially all of them at *λ ≲* 0.5, while the physical rung contains 1. The rungs’ own coverage differs as it should (Fig. 1b,c): the softest rung visits 154 of the 160 cells, the physical rung 79. The total cost is 665 ns of molecular dynamics, 338 ns for each threading including the *λ*_0_ run, against 15.4 *µ*s for the reference.

**Figure 1.**
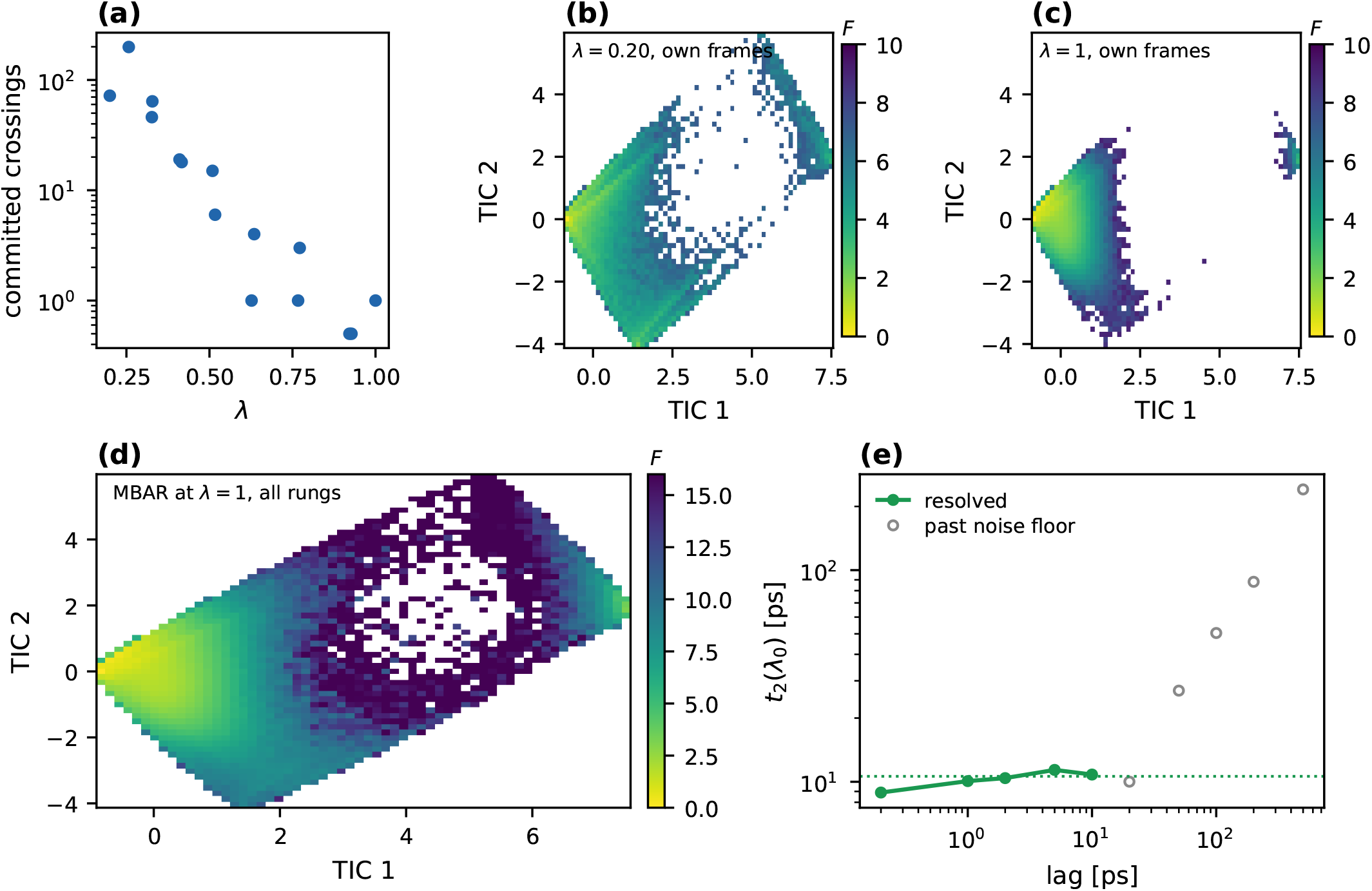
Ladder sampling and MBAR free-energy estimates. **(a)** Committed barrier crossings — a walker reaching one basin core having last been in the other — at each coupling, pooled over the threadings. **(b**,**c)** Free energy in the two leading TICA components from each rung’s *own* frames: the continuous softest rung covers the transition region, while the physical rung stays almost entirely in the reactant basin. **(d)** MBAR at *λ* = 1 from *all* rungs. The transition region is present only because MBAR brings in soft-rung frames with their physical free energy, and the cells the physical rung never visits obtain their stationary weight this way. **(e)** Implied timescale of a direct Markov model on the continuous softest rung against lag. Filled points are resolved (|*µ*_2_| *>* 0.3) and give the internal reference *t*_2_(*λ*_0_) = 10.6 ps, while open points lie past the statistical floor, where the estimate merely tracks the lag.

### 3.3 Thermodynamics from the ladder

#### One threading fails

MBAR on a single threading gave Δ*F* (*B − A*) = 5.50 *k*_B_*T*, 3.7 standard deviations from the reference. The failure is located by a self-consistency test: each rung’s own product-basin fraction divided by the MBAR prediction for its coupling. Rungs up to *λ ≈* 0.41 are in equilibrium: their own product-basin fraction matches MBAR’s prediction for their coupling to within *±*11%. The rung stored while the accumulated weights were non-uniform carries 45% too much product basin, and every rung after the one resampling step carries 30–40% too little: that resample cut the product-basin walkers by two thirds, and 20 ps segments at stiff coupling never cross back to restore them. MBAR draws the physical-rung thermodynamics mainly from those stiff rungs, and inherits the deficit.

#### Variability across ladder threadings

Threading the ladder 4 times from the same continuous run with independent seeds gives product-basin fractions at the physical rung between 0.0029 and 0.0137, and MBAR estimates Δ*F* (*B − A*) = 4.90 *±* 0.54 *k*_B_*T* (range 4.20–5.50). The mean agrees with the reference, but the scatter is 2.6 times the reference’s own uncertainty. The claim that re-sampling “cannot bias the free energy because every frame is retained” is true in expectation and false in realisation, for any basin that a few walkers carry. Pooling threadings reduces it. The pipeline pools 2: seeds 11 and 21, the first two generated, fixed before any downstream result was seen. The other pairs, single threadings and all four are the cost study of Sec. 3.6. In one MBAR problem the two give

|  | two threadings | reference |
| --- | --- | --- |
| $\Delta F(B - A) [k_B T]$ | 5.191 | $4.747 \pm 0.205$ |
| $P(\phi > 0)$ | 0.00628 | 0.0095 |
| barrier A→TS [$k_B T$] | 15.0 | 12.31 |

Two threadings reduce the scatter but do not reach the reference’s own precision: this pair is 5.191 *k*_B_*T*, the six possible pairs give a mean error of 0.28 *k*_B_*T* (at most 0.44), and all four threadings 0.04 *k*_B_*T* (Sec. 3.6). The barrier top of the one-dimensional *F* (*ϕ*) is not recovered either, since it rests on few reweighted transition-region frames. It is not, however, what sets the chain’s rate (Sec. 3.5).

### 3.4 Effects of the TICA basis and cell resolution

The internal reference at the softest rung passes: over lags where the second eigenvalue is resolved, *t*_2_(*λ*_0_) = 10.6 ps with a 13% spread (Fig. 1e), in agreement with the 7.6 ps implied by the committed-crossing rates themselves. At this coupling the product basin is shallow — escape takes ~ 8 ps — while at *λ* = 1 the same process takes 4413 ps. The generator must bridge a factor of 400.

#### A basis at the softest rung places fewer cells on the barrier

With the TICA basis estimated at *λ*_0_ and a 16 *×* 16 tiling, only 10 cells lie on the barrier and 4 at intermediate committor (Table 1). The predicted shift is not affected, but the committor structure the reweighting can act on is thinner: at *λ*_0_ the slowest linear mode is dominated by fast escapes from the shallow basin and by wide soft excursions, not by the barrier geometry relevant at *λ* = 1. Estimating the basis at *λ* = 0.41 raises the barrier cells to 14 at the same 16 *×* 16 resolution.

**Table 1.** Effects of the TICA basis and cell resolution. Each row is a full run of the slow-subspace, cell, MBAR, generator and maximum-caliber stages on the same ladder. *n*_I_ counts cells whose mean torsion lies outside both basin cores, while 0.05 *< q <* 0.95 counts cells at intermediate committor on the prior chain. Prior *t*_2_ is the mean and spread over four seeds at the regulariser selected by validation likelihood. CK counts macrostate elements inside the 95% bootstrap interval of the validation *λ* = 1 data. The last column is the error of the predicted equilibrium shift from the time-resolved density.

| slow subspace and cells | cells | $n_I$ | $0.05 < q < 0.95$ | prior $t_2$ [ps] | $/t_2^{\text{ref}}$ | CK | $\Delta F$ error [$k_B T$] |
| --- | --- | --- | --- | --- | --- | --- | --- |
| $\lambda_0$ basis, $16 \times 16$ | 151 | 10 | 4 | $3722 \pm 57$ | 0.84 | 28/28 | 0.001 |
| $\lambda = 0.41$ basis, $24 \times 16$ | 230 | 23 | – | $8580 \pm 1776$ | 1.94 | 28/28 | – |
| $\lambda = 0.41$ basis, $16 \times 16$ (used) | 160 | 14 | 6 | $5430 \pm 996$ | 1.23 | 28/28 | 0.002 |

#### Refining the cells adds barrier edges no transition constrains

Recovering committor coverage by resolution instead fails at the next stage. A 24 *×* 16 tiling of the same basis places 23 cells on the barrier but has 41 barrier edges, 30 of them with no observed transition at any training lag, against 33 and 22 at 16 *×* 16. The observed transitions on barrier edges fall, because finer cells turn crossings into multi-cell jumps that no single edge records. The prior becomes 1.94 times the reference, with a seed spread of *±*1776 ps (Table 1). The 16 *×* 16 tiling of the *λ* = 0.41 basis is used below: 160 cells, 288 edges, 576 free parameters, a median of 291 frames per cell.

### 3.5 The prior chain

Validation likelihood selects *η* = 10^*−*4^ (NLL 2.4902, against 2.5267 for the next value), and the generator gives *t*_2_ = 5430 *±* 996 ps, a ratio of 1.23 to the reference (Fig. 2). The Chapman–Kolmogorov test places the model inside the bootstrap interval of the validation data in 28 of 28 elements. Using the *λ* = 1 counts directly instead gives *t*_2_ = 2 ps.

**Figure 2.**
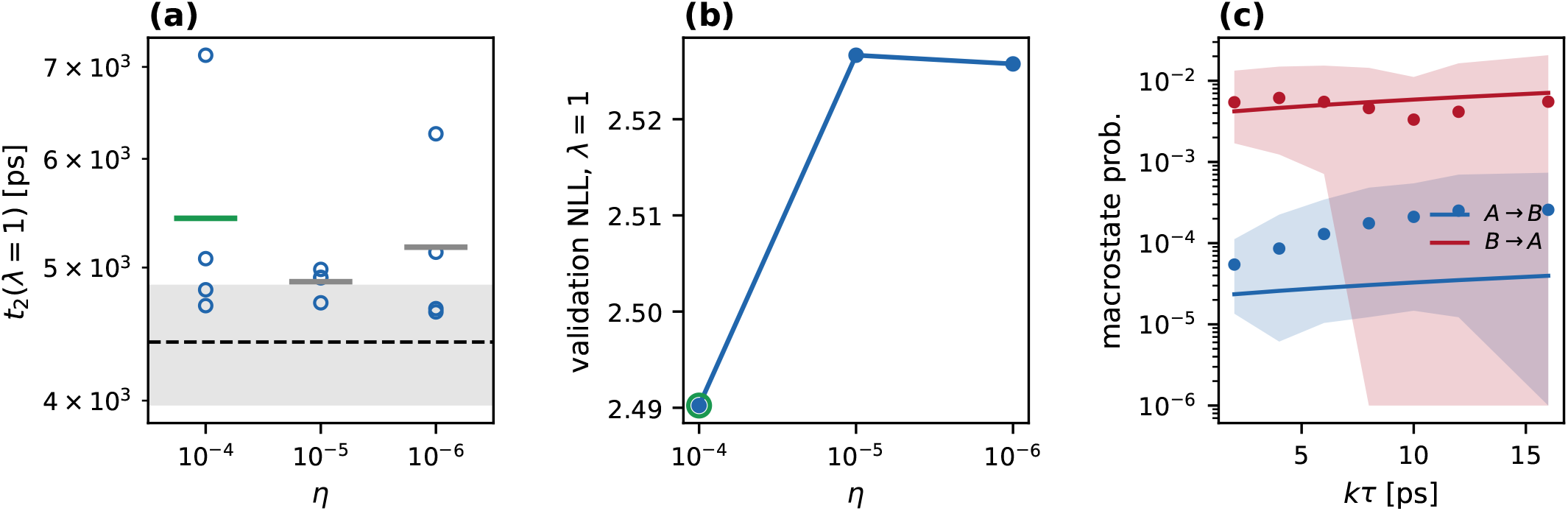
The learned prior at *λ* = 1, transition counts reserved for validation. **(a)** *t*_2_ for four seeds at each regulariser *η*, with bars showing the means and the grey band showing the reference. **(b)** Likelihood of the validation *λ* = 1 counts. The circled value is selected. **(c)** Chapman–Kolmogorov test at *λ* = 1: model lines against the validation data, bands 95% bootstrap over walkers. The trajectories are 20 ps long, so the test reaches 16 ps while *t*_2_ is thousands of ps.

The prior is 1.23 times the reference, with a seed spread of *±* 996 ps. Across all pairs of threadings the ratio ranges from 0.53 to 1.61 (Sec. 3.6), so its deviation is dominated by which threadings were drawn. Three tests locate what does not cause it. *Not the barrier top in π*: a ladder of 2048 *×* 20 ps walkers started from a *λ* = 0.30 pool, with 37 committed crossings, has a far higher MBAR barrier in *F* (*ϕ*) and a faster prior. *Not the initialisation*: kinetic prefactors on barrier edges that no transition constrains do not remain at their initial value but are pulled, through the longer lags, to the same range from initial exponents *−*1, *−*3 and *−*5, and *t*_2_ (3843–6626 ps) varies with the initial value no more than with the seed (Sec. 4). *Not the softest coupling* : removing the continuous *λ*_0_ run from training leaves *t*_2_ at 1.1 times the reference. What the same test does show is where the barrier is constrained: removing every coupling below 0.4 leaves 29 of 33 barrier edges unobserved and makes *t*_2_ 4.5 times the reference.

#### Limits of short-time validation

Across the initialisation tests *t*_2_ ranges from 3843 to 6626 ps while the validation likelihood varies by 0.033, and not in the direction of *t*_2_. The validation *λ* = 1 trajectories are 20 ps long and contain essentially no barrier crossing, so their likelihood, like the Chapman–Kolmogorov test, validates the short-time dynamics and is blind to the barrier rate. Regulariser selection by validation likelihood is therefore a statement about the fast dynamics only, and the slow timescale of the prior is validated solely by the external reference.

#### Learned prefactors and transfer to the physical coupling

The generator does not learn counts at *λ* = 1, but learns one kinetic prefactor per lattice face, and the counts it implies at *λ* = 1 follow from those kinetic prefactors and the exact *π*_1_. Before learning, crossings of the committor surface are observed from 145 of the 160 cells at couplings below 0.6, from 18 between 0.6 and 1, and from 13 on the validation physical rung (Fig. 3a– c). After learning every cell has a crossing probability (Fig. 3d). The information comes almost entirely from the soft couplings. Across the rungs the generator reproduces the counted crossings over four orders of magnitude, from the same starting frames (Fig. 3e). At *λ ≈* 0.92, where none were counted, it predicts about 4 transitions, and on the validation physical rung 4 against 36 observed from a single committed crossing, inside the 95% Poisson interval of that crossing. This agreement is a consistency check, not a test of the barrier rate, which one event cannot provide.

**Figure 3.**
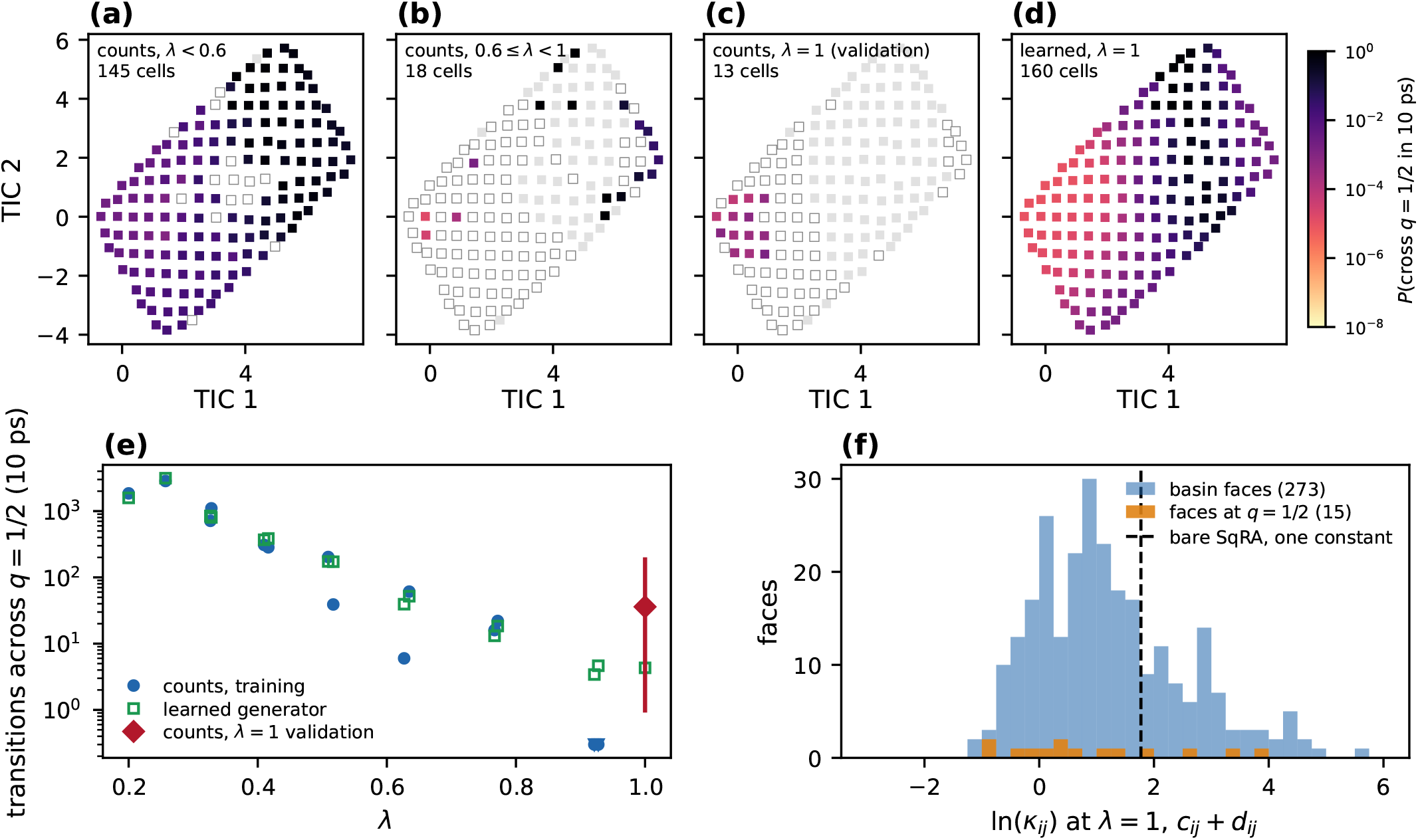
Transfer of learned kinetics to the physical coupling. **(a–c)** Per cell, the fraction of 10 ps transitions that end on the other side of the committor surface 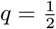, from the counts of the training rungs below *λ* = 0.6, of those between 0.6 and 1, and of the validation physical rung. Grey indicates no frames at those couplings, while open markers indicate frames but no crossing. **(d)** The same quantity from the learned generator at *λ* = 1. **(e)** Transitions across 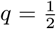 per rung, counted (filled) and predicted by the generator from the same starting frames (open). The diamond marks the validation physical rung, with the 95% Poisson interval of its single committed crossing. **(f)** The learned log-prefactor of every lattice face at *λ* = 1, against the one constant of bare SqRA fitted to the same counts.

The learned slope in the selected regularised fit is negligible where it matters: on the 15 faces that cross 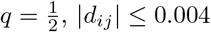. What transfers is therefore a nearly coupling-independent kinetic prefactor per face, measured where crossings are plentiful, combined with the estimated stationary populations at each coupling. That kinetic prefactor is not uniform: across the 288 faces it spans a factor of 106 (5–95%, Fig. 3f). Bare SqRA, which assumes one constant, is the limit *c*_*ij*_ = *c*_0_, *d*_*ij*_ = 0. Fitted to the same training counts and scored by *t*_2_ at *λ* = 1 and the NLL of the same validation counts (one seed each), the nested models give

| exponent | param. | $t_2$ [ps] | NLL |
| --- | --- | --- | --- |
| $c_0$ (SqRA) | 1 | 10980 | 2.5244 |
| $c_0 + \lambda d_0$ | 2 | 1197 | 2.5223 |
| $c_{ij}$ | 288 | 3432 | 2.5123 |
| $c_{ij} + \lambda d_{ij}$ (this work) | 576 | 4817 | 2.4985 |

against 4413 *±* 446 ps. One global rate constant is 2.5 times too slow, or 3.7 times too fast if it may depend on the coupling. One kinetic prefactor per face gives 0.78 of the reference, and its dependence on the coupling the rest. The method’s gain over the constant SqRA baseline used here is the position dependence of the kinetic prefactor, learned from transition counts at the softer couplings.

### 3.6 Cost against brute-force molecular dynamics

The reference MD is itself a convergence experiment. Read as one long run — the first fraction of its independent 30 ns walkers — it gives

| fraction | MD [μs] | $\Delta F(B - A)$ [$k_B T$] | $t_2$ [ps] |
| --- | --- | --- | --- |
| 4.3% | 0.66 | 4.37 | 7604 |
| 8.6% | 1.32 | 4.13 | 9605 |
| 25% | 3.84 | 4.80 | 5767 |
| 50% | 7.68 | 4.86 | 4081 |
| 75% | 11.5 | 4.78 | 4610 |
| 100% | 15.4 | 4.75 | 4430 |

(Fig. 4): Δ*F* settles to within ~ 0.1 *k*_B_*T* of its final value by 25% of the run, *t*_2_ to within ~ 10% only by 50%. Splitting the walkers into disjoint sets gives independent brute-force experiments at a fixed cost: at 0.66 *µ*s, 26% of them contain no committed crossing and their median Δ*F* error is 0.96 *k*_B_*T*. At 1.32 *µ*s it is 0.38 *k*_B_*T*, with *t*_2_ typically off by a factor of 2.2.

**Figure 4.**
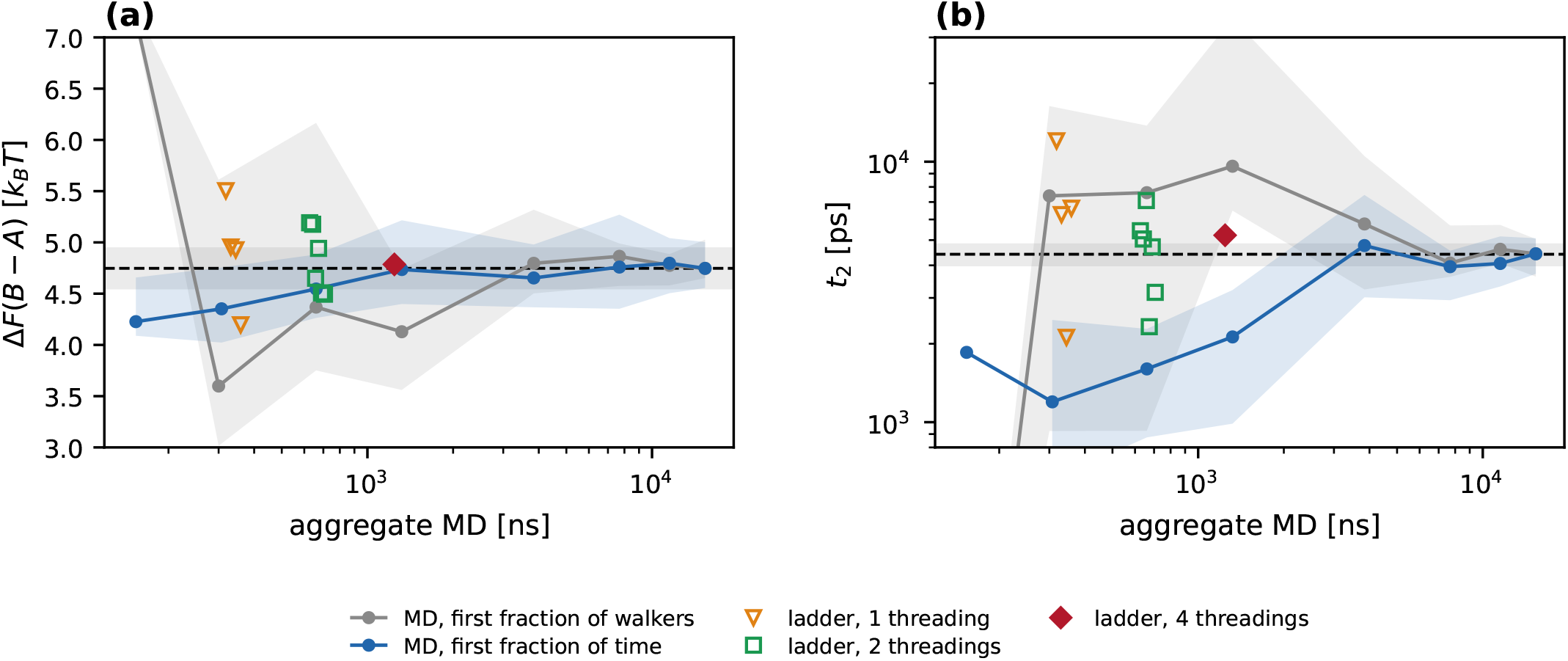
The ladder against brute-force MD at equal cost. **(a)** Δ*F* (*B − A*) and **(b)** *t*_2_ against aggregate MD, the *λ*_0_ run included for the ladder. Lines: the reference MD read as one long run, from the first fraction of its walkers (grey, each 30 ns) or of every walker’s time (blue). The blue curve inherits the equilibrated starting points of all 512 walkers and is optimistic. Bands, 16–84% bootstrap over the walkers used. Symbols: the ladder pipeline (stages 1–5) on each single threading, each pair of threadings, and all four. Dashed line and grey band, the full reference.

The ladder pipeline, run on one, two and four of the threadings from the same continuous *λ*_0_ run, gives (mean absolute Δ*F* error, maximum in brackets)

| thr. | MD [ns] | runs | $\vert \Delta F \text{ err.}\vert $ [$k_B T$] | $t_2/t_2^{\text{ref}}$ |
| --- | --- | --- | --- | --- |
| one | 338 | 4 | 0.42 ($\leq 0.76$) | 0.48–2.73 |
| two | 665 | 6 | 0.28 ($\leq 0.44$) | 0.53–1.61 |
| four | 1321 | 1 | 0.04 | 1.18 |

Two threadings, 665 ns of MD, reach a Δ*F* error that brute force reaches at a few *µ*s, and a typical *t*_2_ error (a factor of 1.31) that it reaches only at ~ 4–8 *µ*s. A single threading is four times cheaper than four, but its prior timescale is off by up to a factor of 2.73 and no internal test flags which threading is the bad one: this is the variance of Sec. 3.3 propagated into the rate. The comparison is for the prior chain. Maximum caliber (Sec. 3.7) run on each single-threading chain recovered the imposed shift to within 0.009 *k*_B_*T* against the density and 0.09 *k*_B_*T* against two measured rates, because it corrects relative to the prior it is given. The accuracy of the final prediction is limited by the prior.

### 3.7 Reweighting with time-resolved density constraints

The target is the prior chain with *k*_*B→A*_ divided by ten, read out on 12 *ϕ*-bins at 14 checkpoints. Its equilibrium consequence is a shift of +2.303 *k*_B_*T* in Δ*F* (*B − A*), withheld from the fit.

#### Conditioning and identifiability of the multiplier basis

With the 28 raw Gaussian bumps the optimum lies on the guard that bounds the coefficients, and moving the guard moves the answer: at |*θ*| *≤* 30 the error in the shift is 0.01 *k*_B_*T* (*χ*^2^ per point 3.25), at |*θ*| *≤* 300 it is 0.08 *k*_B_*T* (1.36), both with the largest coefficient on the bound. The bound, not the caliber, is regularising. Of the 4 coefficients at the bound, 2 act on cells holding ~ 10^*−*6^ of the posterior weight, where rewriting the transitions costs almost no caliber because the caliber is *π*-weighted, and 2 are occupation bumps over *ϕ* and over *q* acting on the same product-basin cells (0.05 of the weight), nearly collinear so that only their combination is determined, a flat valley. Orthonormalising the occupation bumps under ⟨*f, g⟩* = ∑_*i*_ *π*_*i*_*f*_*i*_*g*_*i*_ and the crossing bumps under the equilibrium flux across each committor level removes both: of the 19 occupation directions only 6 carry more than 10^*−*4^ of the leading weighted norm, and all optima become interior. Without a detection floor the largest coefficient is 2.55, unbounded, and a narrower crossing basis (29 directions) changes nothing (0.24 *k*_B_*T*).

#### Sensitivity to the detection floor

With the orthogonal basis and no detection floor the fit is still poor, and 69% of its *χ*^2^ comes from one *ϕ*-bin at the far edge of the product basin, *ϕ ≈* +1.6 rad, where the target density is ~ 10^*−*6^. With log-residuals at uniform *σ* that bin weighs as much as a basin bin, and the tilt distorts the product basin to chase it. No time-resolved measurement resolves such a population. With a detection floor of 10^*−*4^ on the target density the fit and the prediction become

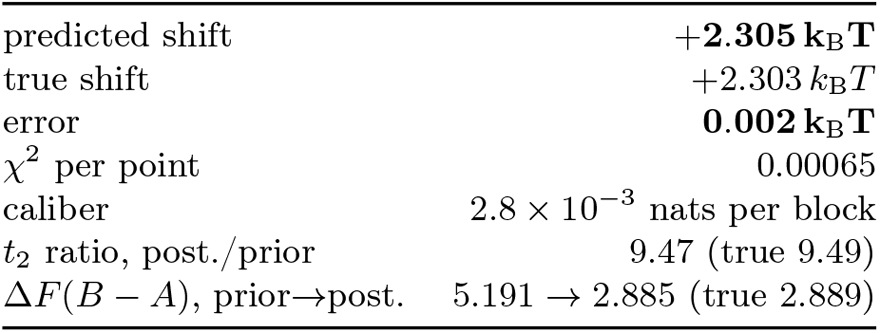

|  |  |
| --- | --- |
| predicted shift | **+2.305 $k_B T$** |
| true shift | +2.303 $k_B T$ |
| error | **0.002 $k_B T$** |
| $\chi^2$ per point | 0.00065 |
| caliber | $2.8 \times 10^{-3}$ nats per block |
| $t_2$ ratio, post./prior | 9.47 (true 9.49) |
| $\Delta F(B - A)$, prior → post. | 5.191 → 2.885 (true 2.889) |

with 15 multipliers, the largest of magnitude 0.3. Nothing thermodynamic was constrained. Because the floor was introduced after the misfit had been localised, its effect is reported in full: the error is 0.007 *k*_B_*T* at a floor of 10^*−*5^ (*χ*^2^ per point 0.0143) and 0.24 *k*_B_*T* at 10^*−*6^ (2.93), while at 10^*−*4^ moving the norm cutoff that discards basis directions from 10^*−*3^ to 10^*−*5^ (12 to 16 directions) leaves the error at 0.004 and 0.004 *k*_B_*T*. The prediction is accurate when the constraint contains only populations a measurement could see, and degrades when it is asked to reproduce populations of cells the identifiable part of the tilt cannot reach.

#### Structure of the inferred free-energy correction

The posterior tracks the withheld truth across the transient and the long-time plateau (Fig. 5b, Fig. 6). The correction Δ*F*_*i*_ is zero in the reactant basin and *−* 2.30 *k*_B_*T* in the product basin (Fig. 5c,d). On the intermediate cells it deepens beyond the product-basin value, to *−* 4.5 *k*_B_*T* on the product side of 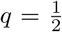— slowing crossings back towards *A* holds population there — but these values are not determined by the data: varying the norm cutoff leaves the fit and the shift unchanged while moving them by up to 8 *k*_B_*T* and the product-basin value by 0.09 *k*_B_*T*. Only the basin-level correction is a result, while the rest lies in the null space of the constraint. Per configuration, Eq. (24) moves *P* (*ϕ >* 0) from 0.0055 to 0.0529.

**Figure 5.**
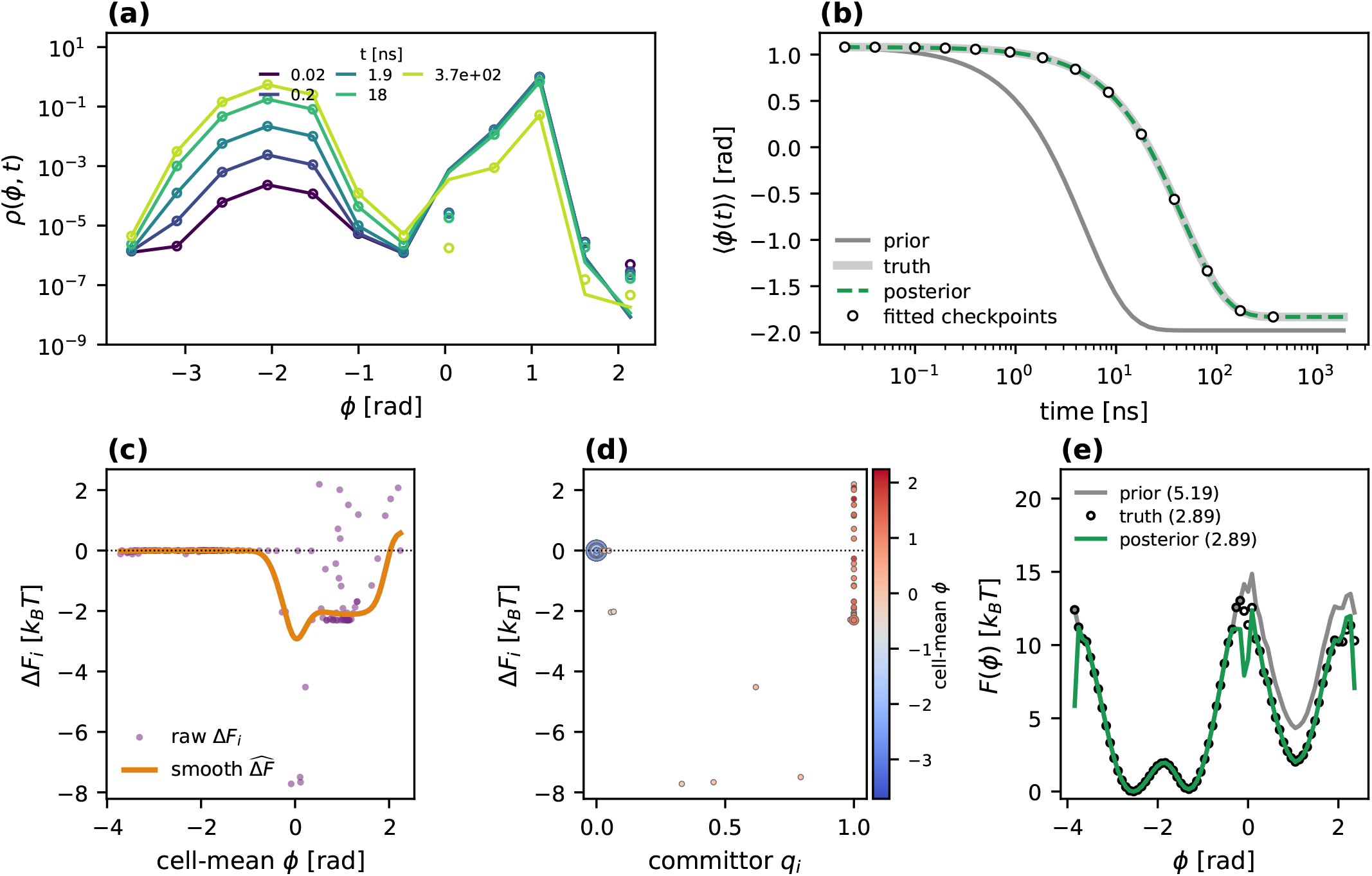
Maximum caliber against a time-resolved density. **(a)** The constraint: circles the target density *ρ*^*\**^(*ϕ*-bin, *t*_*k*_), lines the driven chain at the optimum, 5 of 14 checkpoints labelled in ns. Populations below the detection floor 10^*−*4^ do not enter the objective. **(b) ⟨***ϕ*(*t*) ⟩ relaxing from the product basin, for the prior, the withheld target (*truth*) and the posterior. Circles show the target at the 14 checkpoints the fit sees, while the posterior at every other time, before and after, is predicted by chaining. **(c)** The derived correction Δ*F*_*i*_ = *−*2*k*_B_*T* ln *ψ*_*i*_ against cell-mean *ϕ*, raw and kernel-smoothed. **(d)** The same correction against the committor of each cell, marker area *∝ π*_*i*_, colour the cell-mean *ϕ*. The basin values are determined by the data, while the values on the few intermediate cells are not. **(e)** *F* (*ϕ*) for prior, truth and posterior — a prediction, since *F* enters no objective. Legend values are Δ*F* (*B − A*) in *k*_B_*T*.

**Figure 6.**
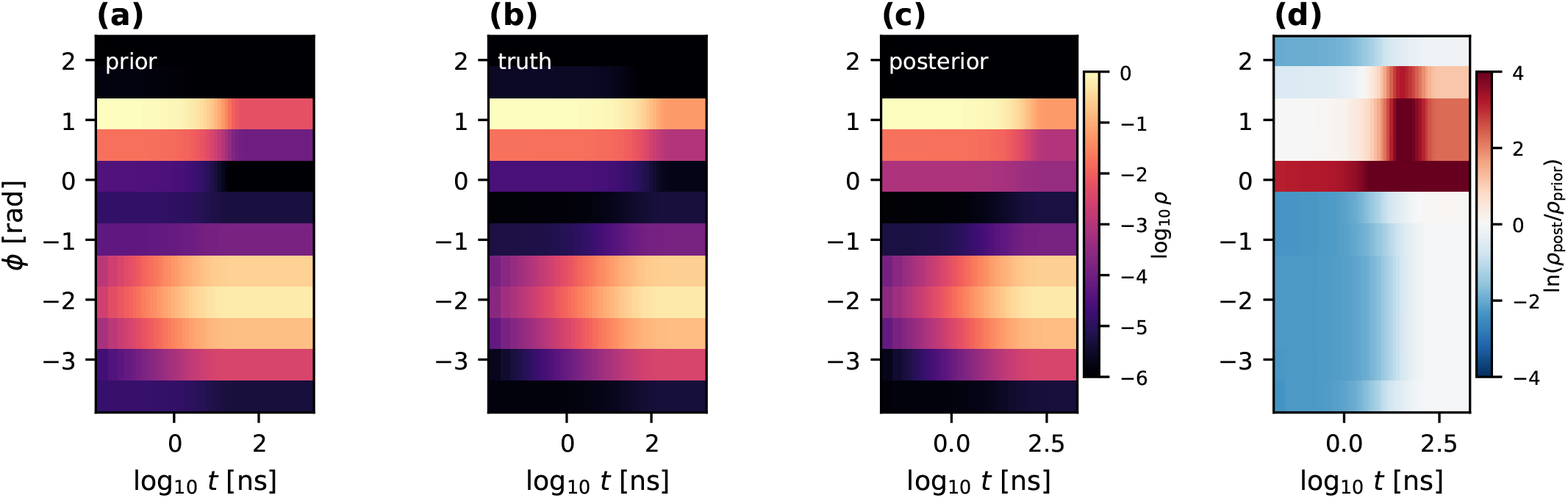
Time-resolved density, prior against posterior. **(a–c)** log_10_ *ρ*(*ϕ, t*) relaxing from the product basin for the prior chain, the withheld target and the maximum-caliber posterior. **(d)** ln(*ρ*_post_*/ρ*_prior_). The largest changes at the barrier bin, *ϕ ≈* 0, lie below the detection floor and are not constrained.

### 3.8 Reweighting with committed-rate constraints

The second experiment constrains rates instead of a density. The target is the prior chain with both committed rates changed: *k*_*B→A*_ made 20 times and *k*_*A→B*_ 3 times faster (*×* 19.83 and *×* 2.99 achieved, with *kτ* = 0.083 at the 20 ps lag). By detailed balance Δ*F* (*B − A*) rises by +1.891 *k*_B_*T*, and *t*_2_ falls from 4818 to 235 ps. The objectives are those of Eqs. (21) and (22): the same tilt and basis as for the density, but two stationary observables in place of 168 transient ones.

#### Two measured rates

With both committed rates constrained the posterior meets them (*×*19.83 and *×*2.99, Fig. 7a) and the shift is +1.892 *k*_B_*T*, an error of 0.000 *k*_B_*T*. This is detailed balance: the ratio of the rates *is* the free-energy difference. Because the committed rates also fix their sum, the posterior’s relaxation time follows, *t*_2_ = 289 ps against 235, and *F* (*ϕ*) matches the truth (Fig. 7b).

**Figure 7.**
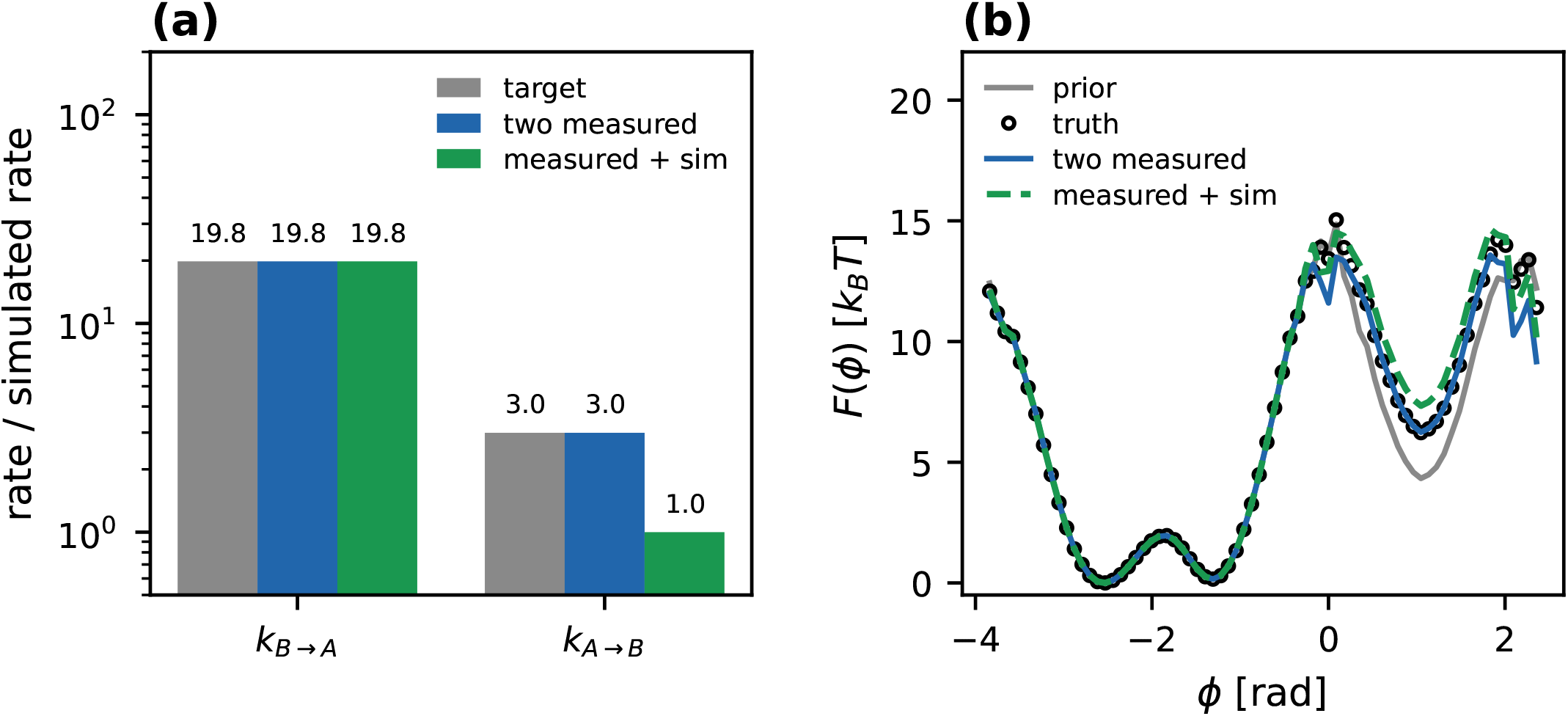
Posterior kinetics and free energies under committed-rate constraints. Target: committed *k*_*B→A*_ made 20 times and *k*_*A→B*_ 3 times faster than in the simulation (the prior chain). *Two measured* : both rates from the target, Eq. (21). *Measured + sim*: *k*_*B→A*_ from the target and *k*_*A→B*_ from the simulation, Eq. (22). **(a)** Committed rates relative to the simulation: the target and the two posteriors. **(b)** The predicted *F* (*ϕ*): with *k*_*A→B*_ from the simulation the product basin comes out too high by the error of that rate.

#### One measured rate and one from the simulation.

When only *k*_*B→A*_ is measured, *k*_*A→B*_ is taken from the prior chain and enters with the same width. Here the simulated *k*_*A→B*_ is wrong by a factor of 3. The fit meets the measured rate and keeps the simulated one (*×* 19.83 and *×*1.00, Fig. 7a), and the shift is +2.988 *k*_B_*T*, equal to the closed form +2.987 and wrong against the truth +1.891 by 1 *k*_B_*T*. That is the error of the simulated rate, 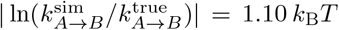, passed one to one into ΔF : the fit does exactly what its assumption asks, and the assumption is what fails: the product basin of *F* (*ϕ*) sits too high by that amount (Fig. 7b). The relaxation time does not show it, *t*_2_ = 292 ps against 289 with both rates measured, because it is set almost entirely by the faster, measured *k*_*B→A*_. A second rate from the simulation is worth exactly as much as the simulation’s rate.

#### One rate alone

Constraining *k*_*B→A*_ with nothing for the other direction meets the constraint and shuts *k*_*A→B*_, to 10^*−*23^ of its prior value, putting the error in the shift at 53 *k*_B_*T*. That completion has *lower* caliber (2.4 *×* 10^*−*5^ nats per block, against 1.7 *×* 10^*−*4^ for the correct one): a rare transition carries almost no path probability, so suppressing it costs almost nothing. The relaxation from the product basin is nevertheless right (*t*_2_ = 287 ps), because it is governed by the measured escape. A single rate determines the kinetics out of its basin and nothing about the equilibrium.

### 3.9 Free-energy corrections in two coordinate projections

Because the posterior reweights the ladder’s own frames, the correction can be drawn in any coordinates (Fig. 8). In the lattice’s own coordinates and in (*ϕ, ψ*) alike, the reactant basin is untouched and the product basin is lowered uniformly. Neither surface enters any objective, and the stored frames carrying the posterior weight are real molecular-dynamics configurations, with no decoder or generative model involved.

**Figure 8.**
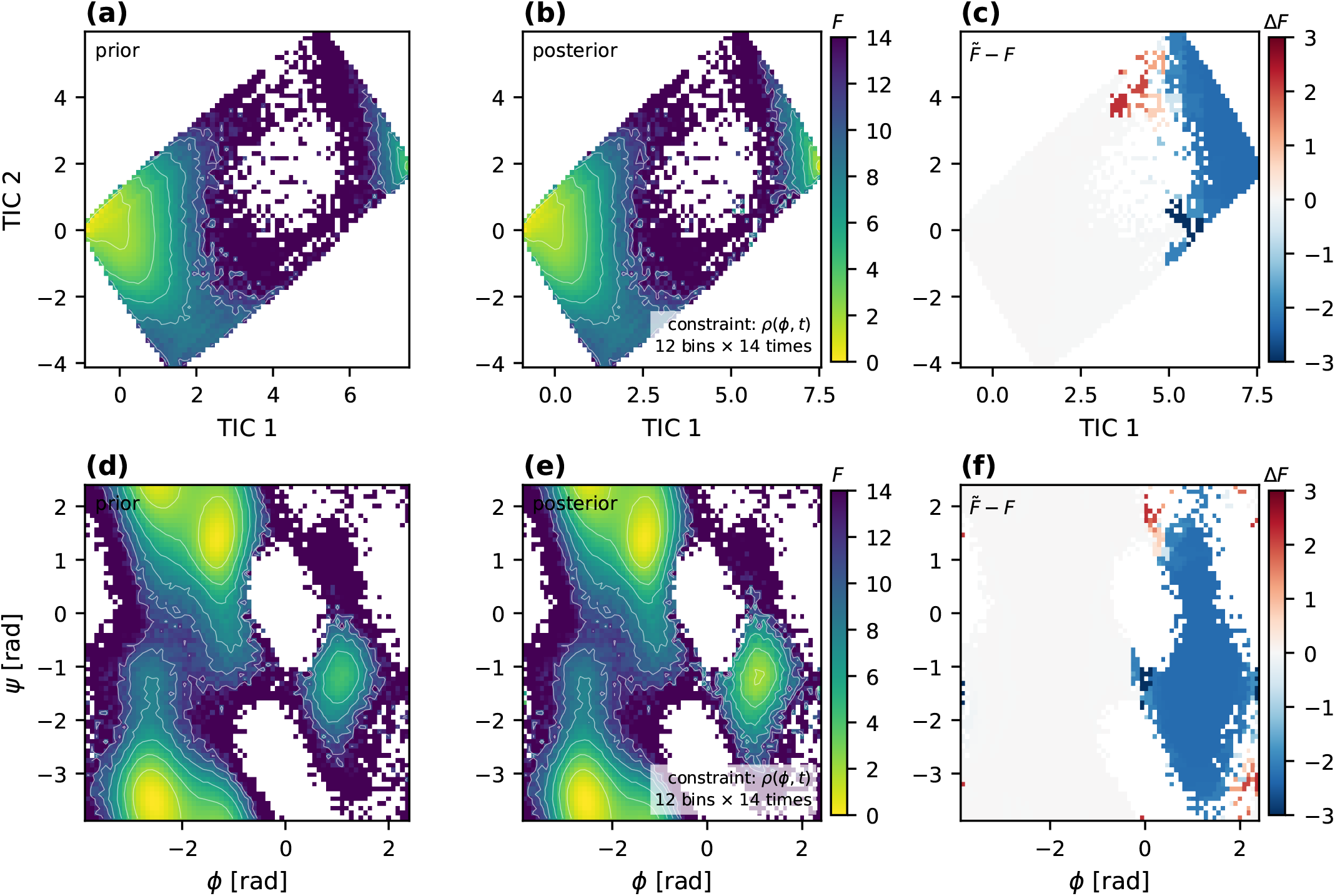
Prior and posterior free-energy surfaces. The posterior is the maximum-caliber chain of Fig. 5, constrained only by the time-resolved density *ρ*(*ϕ, t*) (12 *ϕ*-bins at 14 checkpoints) of a target with *k*_*B→A*_ divided by ten. No rate or free energy is imposed. The top row shows the two leading TICA components in which the cells are defined. The bottom row shows the backbone torsions. Both are per-*frame* surfaces over the same stored configurations, weighted by *W* (*n*) for the prior and *W* (*n*)*ψ*^2^[*c*(*n*)] for the posterior (Eq. (24)), so they differ only through *ψ*. **(c**,**f)** The difference 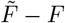 : the correction acts on the product basin and leaves the reactant basin unchanged.

## 4 Discussion

Assembling the method one stage at a time, with an independent check at each, turns “does it work” into “where does it stop working, and why”. Four stages failed on first construction. None of the failures was in the idea that connects them — stationary vector from MBAR, dynamics from the soft couplings, thermodynamics predicted from kinetics — and each located a condition the idea needs.

### The ladder: crossings are set by the coupling

Barrier information per nanosecond is a property of *λ*, not of how the simulation is divided, so the softest rung must be soft enough to cross and still overlap with the next rung. At *λ* = 0.385 a continuous run crosses once every two nanoseconds, while at *λ*_0_ = 0.20 it crosses several times per nanosecond and still reaches the ladder in one FEP step. Threading the rest of the ladder from that run carries its configurations upward, and the soft walker rungs then cross frequently too.

### Thermodynamics: resampling is unbiased only on average

Sequential Monte Carlo resampling is unbiased in expectation, which is the usual justification for treating every stored frame as an equilibrium sample. For a basin carried by a few walkers the expectation is not what one run delivers: one resampling step fixed the product-basin population of every later rung, and a single threading gave Δ*F* with a spread of *±*0.54 *k*_B_*T* over seeds. Pooling independent threadings from one continuous run reduces it, at a cost linear in the number of threadings: two, used here, give a mean error of 0.28 *k*_B_*T*, four 0.04 *k*_B_*T*. Never resampling does not remove it: four threadings that carry the accumulated weights into a weighted MBAR give Δ*F* = 4.86 *±* 0.57 *k*_B_*T* one at a time, against *±* 0.54 with resampling, and 4.750 *k*_B_*T* pooled. Without resampling the weights degenerate instead, and the product basin is again carried by a few walkers. The spread is the sampling error of a rare basin, and independent threadings reduce it as 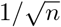, so they buy an error bar, not a correction.

### Prior kinetics and limits of validation

At the physical coupling the ladder contains 1 committed crossing, and most barrier edges of the cell chain carry no observed transition at any training lag. Their rates are nonetheless not arbitrary: the *π*_*λ*_ factor fixes their thermodynamics, and the longer lags pull their kinetic prefactors to a common range from any initialisation. What constrains the barrier is the soft couplings: removing *λ <* 0.4 from training leaves 29 of the barrier edges unobserved and makes *t*_2_ 4.5 times the reference. Two limits follow. The prior’s slow timescale is 1.23 times the reference for the pair of threadings used, 0.53–1.61 across all pairs and 1.18 with all four. The barrier top of MBAR, the initialisation and the softest coupling are each excluded as causes, and what remains is dominated by the sampling of the threadings. And no internal test at *λ* = 1 sees the slow timescale: the validation likelihood and the Chapman–Kolmogorov test are computed on 20 ps trajectories without crossings, and the likelihood varies by 0.033 while *t*_2_ ranges from 3843 to 6626 ps. The fast dynamics are validated internally, while the barrier rate is validated only externally.

### Limitations and future work: kinetic prefactor

The log-linear assumption neglects curvature, which can affect transferred rates while leaving the imposed stationary populations unchanged. A possible extension would be

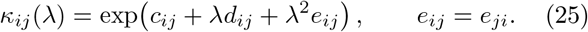

This form preserves positivity and detailed balance in the untruncated model, and reduces to the present model when *e*_*ij*_ = 0. The additional curvature coefficients would require regularisation and validation on couplings not used for fitting or regulariser selection to assess whether they improve predictive accuracy. This extension is left for future work. All results reported here use the log-linear prefactor.

### The cells: coverage and constraint pull in opposite directions

The reweighting acts through the committor, so it needs cells across the barrier, while the generator needs observed transitions on the edges between them. Refining the tiling raises the first and lowers the second, because finer cells turn observed crossings into multi-cell jumps no single edge records, and at 24 *×* 16 the unconstrained barrier edges make the prior 1.94 times the reference and seed-unstable. The basis rung decides how much coverage a given resolution buys: a basis estimated at *λ*_0_, where the slowest linear mode is the shallow basin’s fast escape, places 4 cells at intermediate committor, and one at *λ* = 0.41 places 14 on the barrier and 6 at intermediate committor at the same resolution. For the predicted Δ*F* this did not matter once the objective was well posed (Table 1), but it matters for whether the correction has any committor structure to show.

### The reweighting: identifiability and what a measurement can see

The maximum-caliber stage of the parent construction assumed that the caliber is the only regulariser. It is not, when the multiplier basis has directions acting on cells of negligible posterior weight, or collinear directions: the caliber is *π*-weighted and does not penalise the first, is flat along the second, and a bound on the coefficients silently takes its place. A *π*-weighted orthogonal basis restores the premise, and exposes the remaining issue: an objective that weighs unmeasurable populations as heavily as observable ones. The prediction is accurate with a detection floor a measurement could plausibly have, and not below it. The same analysis shows that only the basin-level correction is determined, while per-cell corrections on the barrier lie in the null space of the constraint.

### Rates: what a single measurement can and cannot say

Detailed balance makes a rate pair a free energy and leaves a single rate without thermodynamic content, and maximum caliber does not fill the gap neutrally: suppressing a rare transition costs almost no caliber, so the least-committal completion of one measured rate closes the barrier in the other direction. The parent construction’s single-rate error of 1.45 *k*_B_*T* was set by a bound on its coefficients rather than by the physics. With an identifiable basis the same chain gives 4.18 *k*_B_*T*. Adding the relaxation time as a second constraint does not repair it and can make it far worse, because after the perturbation the slowest mode of the chain need not be the basin exchange. What does is an explicit statement of where the unmeasured rate comes from — the simulation, with its uncertainty — after which Δ*F* is exactly as accurate as that rate: a simulated rate off by a factor of 3 puts 1 *k*_B_*T* into Δ*F*. Committed rates, with the committor recomputed on the posterior chain, fix the sum of the two rates as well as their ratio, so two measured rates also give the relaxation time.

### Cost

The pipeline is 665 ns of molecular dynamics, 338 ns per threading. Brute-force MD reaches the same Δ*F* accuracy only at a few *µ*s and a comparable *t*_2_ at ~ 4–8 *µ*s, and at the pipeline’s own cost a quarter of brute-force experiments contain no barrier crossing at all (Sec. 3.6). The number of threadings is the lever: one is cheaper but leaves the prior timescale uncertain by up to a factor of 2.73, four bring Δ*F* to 0.04 *k*_B_*T*. The generator fit takes minutes on one GPU and the maximum-caliber fits tens of minutes on CPU.

## 5 Conclusions

We have introduced Anneal-MaxCal, an approach that determines the states, populations and interconversion rates of a molecular system from an annealed solute-tempering ladder, and corrects them against kinetic data by maximum caliber. MBAR provides the equilibrium populations at every coupling, and a square-root-approximation generator, whose position- and coupling-dependent kinetic prefactors are learned from transitions at the softer couplings, transfers the dynamics to the physical Hamiltonian. The transition counts of the physical rung are used only for validation, and no predefined reaction coordinate is required. For alanine dipeptide, the prior model yields a slowest relaxation time of 5430 *±* 996 ps, compared with 4413 *±* 446 ps from 15.4 *µ*s of unbiased dynamics. It does so from 20 ps trajectories, 665 ns of simulation in total and a single committed barrier crossing at the physical coupling, an accuracy that brute-force simulation reaches only after several *µ*s.

Maximum-caliber reweighting of this prior, with the posterior obtained by a generalised Doob transform, converts kinetic constraints into thermodynamic predictions, since no thermodynamic quantity enters any objective. A synthetic time-resolved density corresponding to a tenfold slower back-transition predicts the resulting free-energy shift to within 0.002 *k*_B_*T*. A pair of forward and backward rates recovers the shift to within 10^*−*3^ *k*_B_*T* and gives a relaxation time of 289 ps against 235 ps. A single rate does not determine the equilibrium. Maximum caliber completes it by suppressing the unmeasured transition, and when that rate is instead taken from simulation, its error propagates directly into the free energy.

Anneal-MaxCal requires the softest coupling experience barrier crossings while retaining overlap with the ladder. The discretisation must cover the transition region without leaving barrier edges unconstrained by observed transitions. The reweighting basis must be identifiable against a forward model reporting on kinetics. Extending the approach to larger systems, to multi-state landscapes and to experimental kinetic data, together with a nonlinear coupling dependence of the kinetic prefactors, will be the subject of future work.

